# Skin microbiome composite features in Atopic Dermatitis via integration analysis across cohorts

**DOI:** 10.1101/2025.05.18.653690

**Authors:** Yihua Wang, Bingqiang Liu

## Abstract

Disruption of the skin microbiota is closely associated with the onset and progression of Atopic Dermatitis (AD). However, inconsistencies across studies have hindered a comprehensive understanding of their role in AD and their potential as reliable diagnostic biomarkers. To address this, we conducted a cross-cohort integrative analysis (CCIA) of raw 16S rRNA sequencing data and metadata from the largest available dataset to date, encompassing 1,522 samples across 10 independent studies. We identified consistent microbial signatures distinguishing AD patients from healthy controls. Significant alterations were observed in both α-diversity and community composition between AD and control groups, while lesional and non-lesional sites within AD patients showed no significant differences. Given the impact of confounding factors on the skin microbiota, we applied MMUPHin framework to correct for batch effects and performed subgroup analyses based on different batches. Differential taxa were identified using Permutation testing, Wilcoxon rank-sum tests, and LEfSe analysis. These features were used to develop predictive models with four machine learning algorithms, achieving high diagnostic accuracy in an independent validation cohort (AUROC = 0.83). Our study provides a comprehensive reference of skin microbial alterations in AD, offering valuable insights into host–microbe interactions and highlighting their potential as diagnostic biomarkers for early detection and targeted therapeutic strategies.

## INTRODUCTION

Recent studies have increasingly implicated alterations in the skin microbiota are associated with pathogenesis of various human diseases, notably atopic dermatitis (AD) ^1–3^. AD, a prevalent chronic inflammatory skin disorder with rising global incidence, has emerged as a significant public health concern^4^. While traditionally more common in developed nations, its prevalence is now increasing in developing countries^5,6^. Typically manifesting in early childhood, AD often persists into adulthood^7,8^, significantly impacting the quality of life and underscoring the importance of early diagnosis and intervention^9–11^. The pathogenesis of AD is multifactorial, involving a complex interplay of genetic predisposition, environmental exposures, and immune dysregulation. These include epidermal barrier dysfunction, exaggerated immune responses characterized by increased allergen penetration and IgE sensitization, and dysbiosis of the skin microbiota^12,13^.

Recent advances in methodologies and technologies for studying skin-associated bacteria have enabled investigations at the strain level, providing deeper insights into the relationship between alterations in the skin microbiota and disease^14^. Emerging evidence suggests that changes in the composition and function of the skin microbiome are linked to the pathogenesis of AD^13,15^. Unlike the acute perturbations caused by viral or bacterial pathogens, commensal microbial communities can more subtly modulate immune responses and contribute to epidermal barrier dysfunction—hallmarks of AD pathophysiology^16,17^. Multiple studies have demonstrated that specific microbial taxa, particularly *Staphylococcus aureus*, tend to dominate the skin microbiota of individuals with AD^18,19^. *S. aureus* has been isolated from both lesional and non-lesional skin of AD patients ^20,21^. The reduced microbial diversity in AD skin promotes the colonization of *Staphylococcus*, particularly *S. aureus*, within lesional sites^6,22,23^. The expression of numerous virulence factors by *S. aureus*, combined with the diminished production of antimicrobial peptides in AD skin, collectively contributes to disease pathogenesis^24,25^. A strong correlation has been observed between disease severity and reduced skin bacterial diversity, particularly in areas predisposed to AD^22^. Modulating microbial diversity has been proposed as a potential therapeutic strategy for AD^26^.

16S rRNA sequencing provides a powerful approach to identify disease-associated microbes and explore underlying pathogenic mechanisms. Although previous studies have reported certain characteristics of the skin microbiome^13,27,28^, discrepancies between studies present challenges in validating these features across different populations, thereby limiting the diagnostic potential of microbiome-based markers in AD^27,29–32^. Whether differences in skin microbiome composition and structure can serve as reliable microbial biomarkers for AD remains an open question. Additionally, variations in amplicon sequencing analyses across studies—including differences in geographic regions, sequencing platforms, and 16S rRNA gene target regions—further complicate efforts to achieve standardized and systematic analyses. Therefore, there is an urgent need for multinational, large-scale cohort studies, standardized sampling and analytical methods, and model systems to identify consistent microbiome signatures associated with AD. Cross-cohort integrative analysis (CCIA) is expected to overcome these challenges by evaluating the robustness of disease-microbiome associations through the comparison of multiple case-control studies. The primary goal of CCIA is to identify consistent associations across cohorts, thereby minimizing the impact of biological and technical confounders. CCIA has demonstrated strong performance in previous studies, highlighting its potential as a valuable tool across various research fields^33–35^.

In this study, we integrated raw 16S rRNA sequencing data and associated metadata from 10 publicly available datasets, encompassing a total of 1,522 skin samples, to perform a cross-cohort integrative analysis (CCIA) using a standardized bioinformatics pipeline. Our aim was to identify consistent features of the commensal skin microbiome. Given the influence of confounding factors on skin microbiota, we applied MMUPHin^36^ to correct for batch effects and conducted subgroup analyses across different cohorts. To identify microbial composite biomarkers associated with atopic dermatitis (AD), we applied a combination of statistical approaches, including permutation tests^37^, Wilcoxon rank-sum tests^38^, and linear discriminant analysis effect size (LEfSe)^39^. Based on the resulting microbial signatures, we constructed diagnostic models using four machine learning algorithms to evaluate their potential in improving the accuracy of AD detection. Collectively, our study has the potential to provide novel insights and valuable resources for investigating host-microbiome interactions, early diagnosis, and targeted interventions in AD.

## RESULTS

### Workflow for cross-cohort integration analysis of skin metagenomics in AD

In this study, we conducted a systematic search and screening of studies from the PubMed database, as outlined in the flowchart (Fig. 1), identifying a total of 149 studies.

**Fig. 1.**
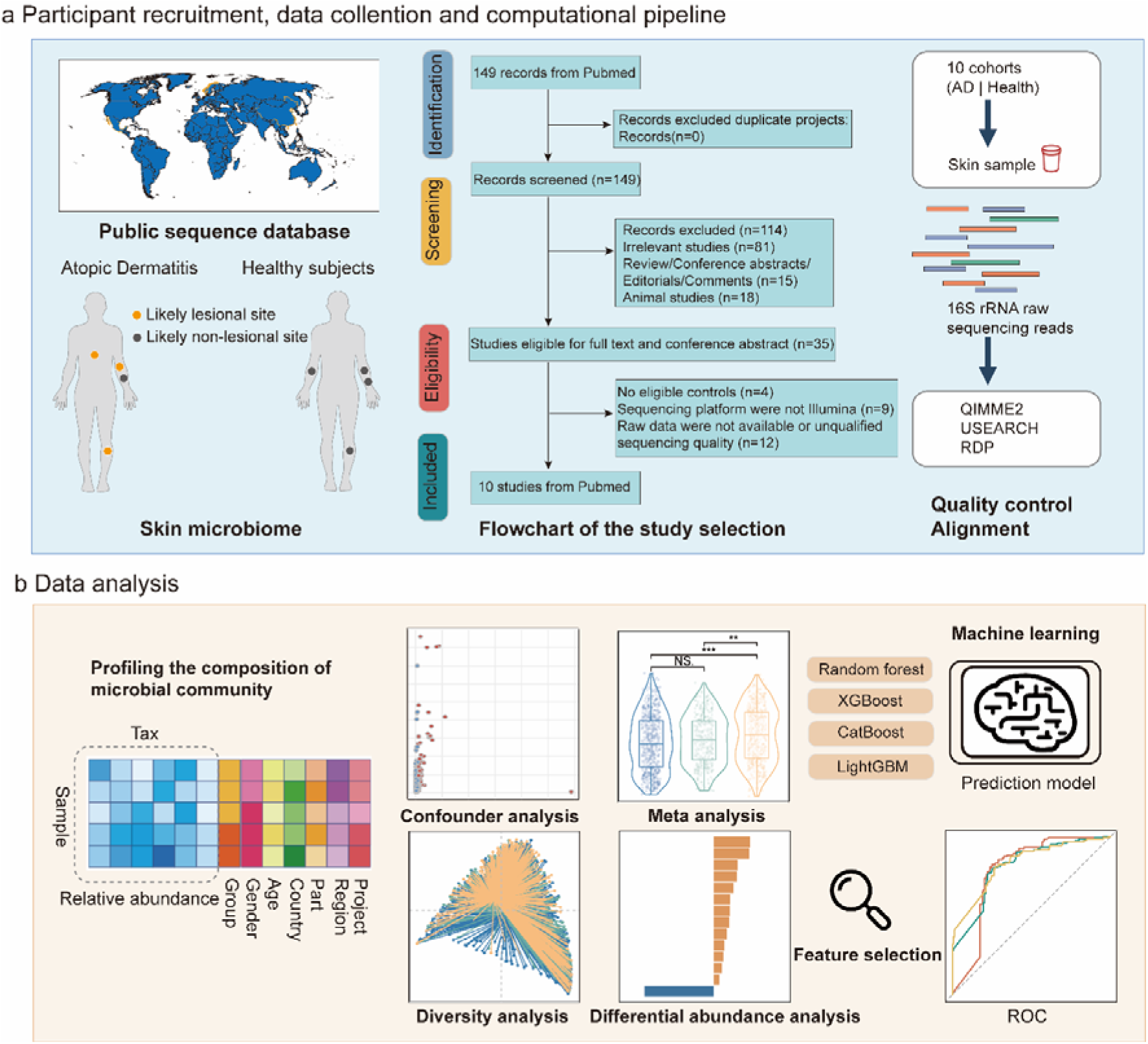
Workflow for cross-cohort integration analysis of skin microbiome in AD. **a** This study included 10 skin microbiome 16S rRNA gene sequencing cohorts (n = 1522) from Atopic Dermatitis (AD) studies across different global geographic regions. Data retrieval and study selection were performed based on a meta-analysis search strategy in the PubMed database. All raw sequencing data were processed using QIIME2 and USEARCH, with taxonomic classification annotated through the RDP database. **b** Detailed bioinformatics analysis was conducted based on the microbiome abundance data and associated metadata. This included analyses of confounding factors, random effects models, diversity analysis, and feature selection. Machine learning models were subsequently constructed, leveraging these features for the diagnosis of Atopic Dermatitis.

After an initial screening of titles and abstracts, we excluded reviews, animal studies, and studies focusing on the gut microbiome or other unrelated topics, identifying 35 studies for full-text assessment. Ultimately, 16S rRNA gene sequencing data from 10 studies on the skin microbiome in atopic dermatitis (AD) met the criteria for further analysis. These 10 datasets were designated as S1 to S10^13,40–48^. This integrated dataset includes 726 participants (419 AD patients and 307 healthy controls), comprising a total of 1,522 samples. Samples were categorized into three groups: AD lesional sites (748 samples), non-lesional sites (435 samples), and healthy controls (339 samples). The sample size of individual studies ranged from 7 to 111 participants. Notably, studies S4, S5, and S6 were not case-control studies and included only AD patients without healthy controls. Due to unavailable age information in some studies, the average age of participants remains unclear. All studies employed Illumina sequencing platforms. With the exception of S1 and S6, which targeted the V1–V3 region, all other studies sequenced the V3–V4 region of the 16S rRNA gene. The included studies were conducted in China, Denmark, Mexico, Norway, Germany, the Netherlands, the United States, Switzerland, and Poland. Table 1 provides detailed information on the demographic characteristics, hypervariable regions, and sequencing platforms of the included studies. To ensure consistency in bioinformatics analysis, all raw sequencing data were reprocessed using USEARCH and QIIME2 (Fig. 1b). Furthermore, our study aimed to uncover patterns of skin microbiome alterations through comprehensive statistical analyses and to leverage machine learning algorithms for AD diagnosis. We first conducted a confounding factor analysis to assess the impact of different variables on sample grouping. Next, we identified 29 AD-associated microbial biomarkers through a series of differential analyses and feature selection approaches, both across cohorts and within sub-cohorts. Classification models based on Random Forest, XGBoost, CatBoost, and LightGBM algorithms were trained on the discovery cohort and subsequently evaluated on an independent validation cohort. The predictive performance of these models demonstrated that the identified microbial biomarkers can effectively distinguish individuals with atopic dermatitis (AD), highlighting their potential utility in microbiome-based diagnostics.

**TABLE 1.**
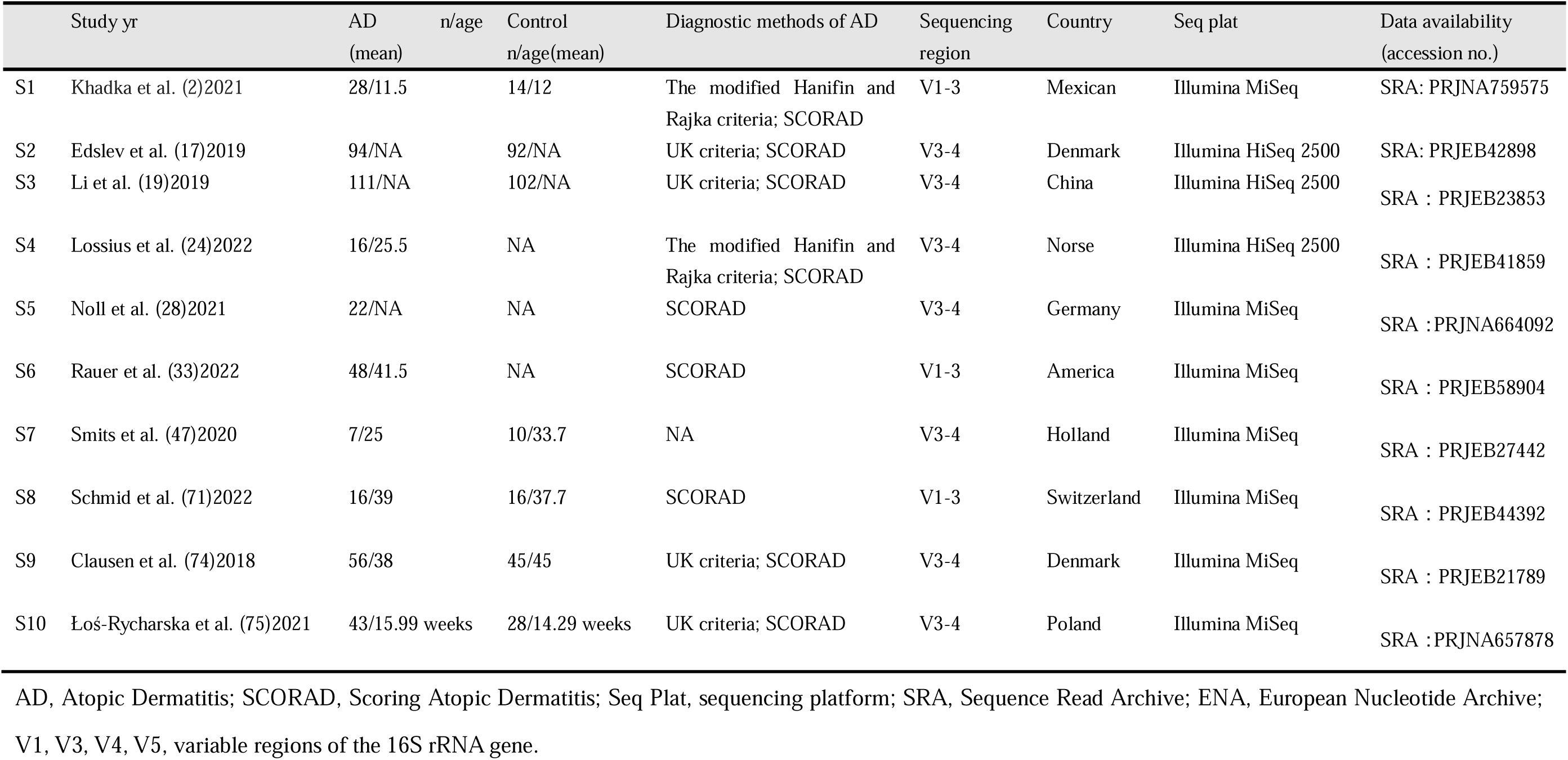
Characteristics of included projects in the analysis

### Assessment and correction of confounding factors in cross-cohort AD microbiome analysis

Following data acquisition, samples were categorized into three groups: atopic dermatitis lesional sites (ADLS), non-lesional sites (ADNLS), and healthy controls (HC). Given the heterogeneity across the ten included datasets—including differences in sequencing batches, 16S rRNA gene target regions, geographic origins, and sampling sites—we systematically evaluated the impact of these four factors on microbiome composition and sample classification. To assess potential confounding effects, we conducted an ANOVA-based confounder analysis, quantifying the proportion of microbial abundance variance attributable to AD status relative to that explained by each confounder. The analysis revealed that alterations in the abundance of specific microbial taxa were, in some cases, more strongly influenced by technical or geographical factors than by disease status. Notably, inter-cohort variation in the relative abundance of certain taxa frequently exceeded the differences observed between AD and control samples (Fig. 2a–d), underscoring the need for rigorous correction of confounding effects in cross-cohort microbiome studies.

**Fig. 2.**
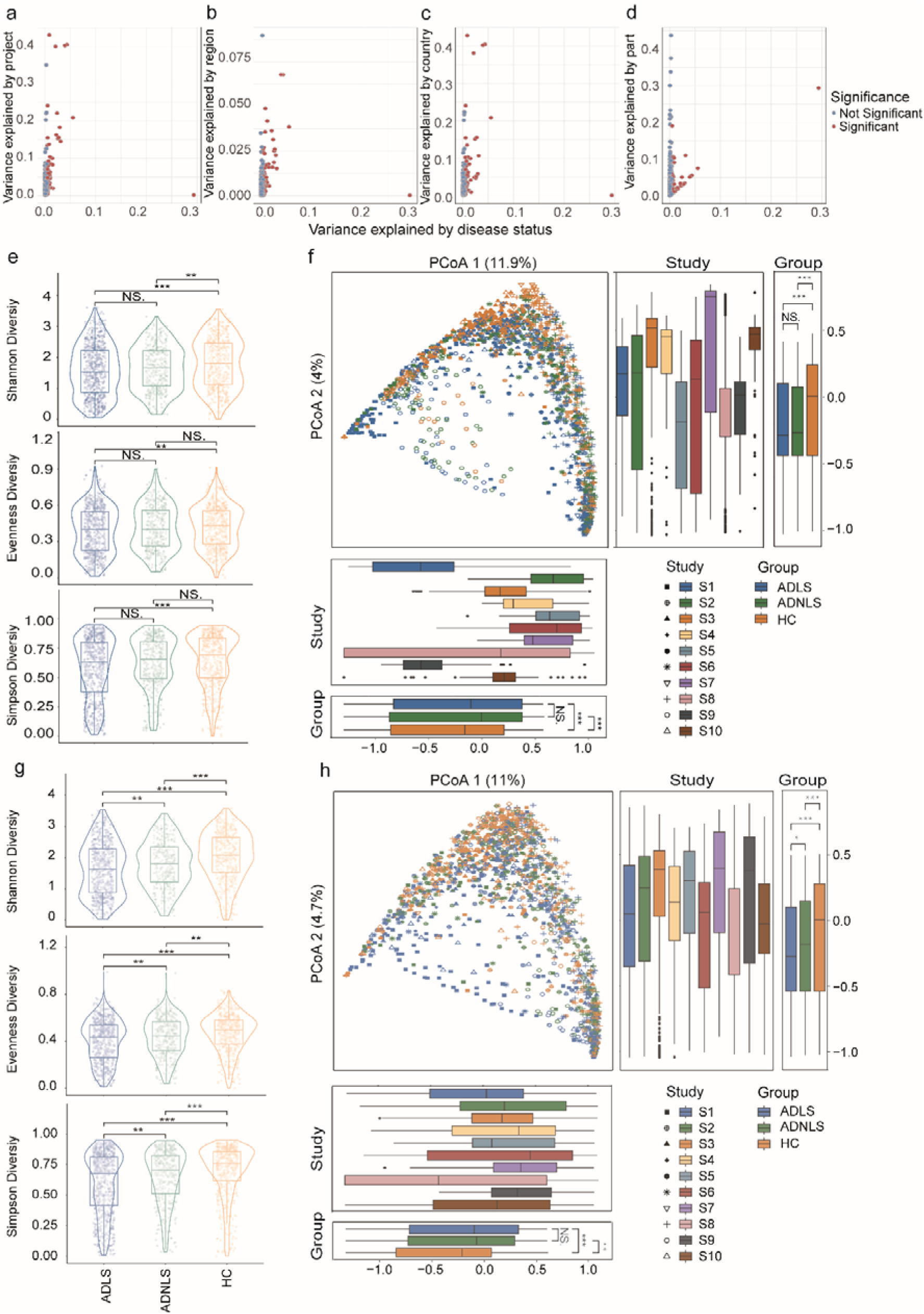
Confounder analysis and removing batch effect. **a-d** This analysis aims to investigate the relationship between the variance explained by disease status (AD vs. control) and various potential confounding factors for each taxa, including cohort differences, sequencing regions, countries, and sampling sites. Among the microbes analyzed, 94 exhibited significant differences between AD and control groups, highlighted in red. All available samples were considered to estimate the variance explained by each confounding factor. **e, g** Display the α-diversity of ADLS, ADNLS, and HC before and after batch effect correction, measured by the Shannon index, Evenness index, and the Simpson index. Batch effects were removed using the MMUPHin tool. Differences in α-diversity between groups were assessed using the Wilcoxon test. “***” denotes p < 0.001, “**” denotes p < 0.01, “*” denotes p < 0.05, and “NS” indicates no significant difference between groups. **f, h** Show the results of principal coordinate analysis (PCoA) before and after batch effect correction using MMUPHin. PCoA results reveal significant differences in microbial composition among different disease states (P = 0.001) and across cohorts (P = 0.001). The significance of β-diversity based on Bray-Curtis distances was calculated using the PERMANOVA method with 999 permutations (two-tailed). Data in the box plots are presented as interquartile ranges (IQRs), with the median indicated by a black horizontal line, and whiskers extending to the farthest data points within 1.5 times the IQR.

To mitigate batch effects across the 10 cohorts, we applied MMUPHin (Meta-Analysis Methods with a Uniform Pipeline for Heterogeneity in microbiome studies). We then conducted a permutational multivariate analysis of variance (PERMANOVA) based on Bray–Curtis dissimilarity to evaluate the extent and significance of cohort and disease status effects on microbiome structure before and after batch correction. Prior to correction, cohort differences accounted for 26.64% of the variance in microbial community structure (PERMANOVA, R² = 0.2664, P ≤ 0.001), which was reduced to 6.89% after correction (PERMANOVA, R² = 0.0689, P ≤ 0.001) (Supplementary Table 1-1). Principal coordinate analysis (PCoA) further confirmed that differences between cohorts along PCoA1 and PCoA2 were reduced after correction, although cohort effects on microbial community structure remained statistically significant (Fig. 2f, h).

Additionally, batch correction substantially reduced the influence of confounding factors such as sequencing region (PERMANOVA, pre-correction: R² = 0.028, F = 41.133, *P* ≤ 0.001; post-correction: R² = 0.007, F = 10.205, *P* ≤ 0.001) and country (PERMANOVA, pre-correction: R² = 0.228, F = 53.207, *P* ≤ 0.001; post-correction: R² = 0.066, F = 12.690, *P* ≤ 0.001), with the variance explained decreasing from 2.8% to 0.7% and from 22.8% to 6.6%, respectively. In contrast, the differences in microbiome composition between AD lesional, non-lesional, and control groups remained significant before and after correction, with minimal impact on R² and F values (PERMANOVA, pre-correction: R² = 0.028, F = 20.691, *P* ≤ 0.001; post-correction: R² = 0.026, F = 19.418, *P* ≤ 0.001) (Supplementary Table 1-1). Given these findings, all subsequent analyses of overall microbial community structure across cohorts were performed using the batch-corrected data.

### Alpha diversity analysis between AD lesion, non-lesion, and control groups

To investigate whether alterations in skin microbial richness and diversity are associated with AD, we performed alpha diversity analyses. The Shannon diversity index (Permutation, ADLS|HC:P < 0.001; ADLS|ADNLS: P < 0.01), Pielou’s evenness index (Permutation, ADLS|HC: P < 0.001; ADLS|ADNLS: P < 0.01), and Simpson diversity index (Permutation, ADLS|HC: P < 0.001 ; ADLS|ADNLS: P < 0.01 ) all indicated significant differences in microbial richness and evenness between the AD and control groups, with greater divergence observed between AD and control groups than between lesion and non-lesion AD sites.

Although MMUPHin was used to mitigate batch effects, potential confounding factors affecting the microbiome composition remain. Therefore, we conducted study-specific subgroup analyses. Across the ten included studies, alpha diversity in the ADLS group was significantly lower than or comparable to that in the healthy control group (Fig. 3a, c, e). A random-effects model was selected for meta-analysis, which revealed significant differences between the ADLS and healthy control groups in the Shannon diversity index (SMD = -0.35, 95% CI = [-0.67, -0.03], *P* = 0.0303), Pielou’s evenness index (SMD = -0.09, 95% CI = [-0.16, -0.02], *P* = 0.0137), and Simpson diversity index (SMD = -0.1, 95% CI = [-0.2, -0.01], *P* = 0.0349) (Fig. 3b, d, f).

**Fig. 3.**
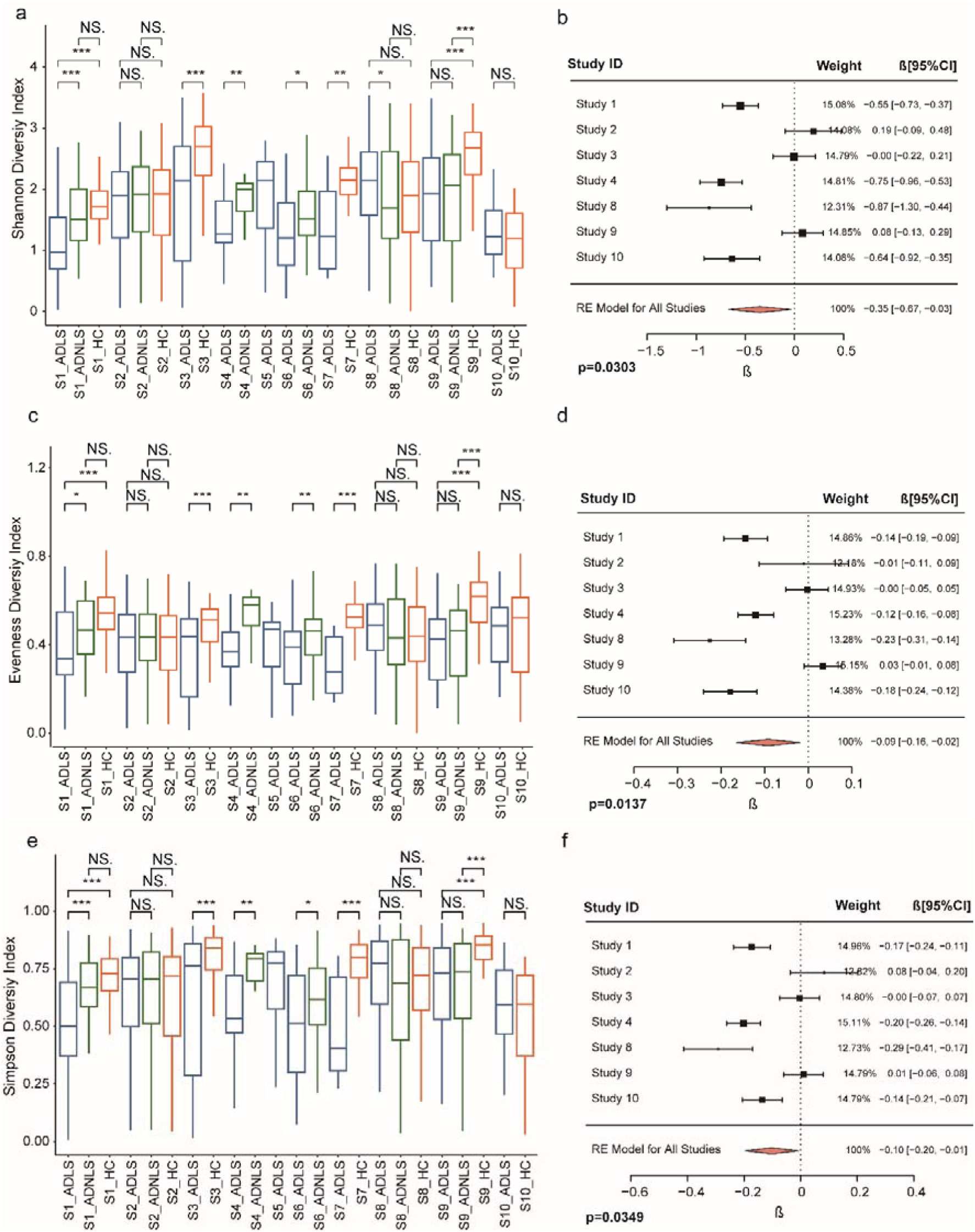
α diversity analysis. **a, c, e** Comparison of α-diversity between the ADLS, ADNLS, and HC groups in different studies, measured using the Shannon diversity index (a), Evenness diversity index (c), and Simpson diversity index (e). **b, d, f** Meta-analysis of α-diversity between the ADLS and HC groups, with standardized mean differences (SMD) and 95% confidence intervals (CIs) for the Shannon diversity index (b), Evenness diversity index (d), and Simpson diversity index (f) presented in forest plots. Statistical significance is indicated as “***” denotes p < 0.001, “**” denotes p < 0.01, “*” denotes p < 0.05, and “NS” indicates no significant difference between groups.

### Characterization of AD-associated microbiome composition

To assess compositional differences in the skin microbiome across disease states, we performed β-diversity analyses using Bray–Curtis and Jaccard dissimilarity metrics. Principal coordinate analysis (PCoA) based on Bray–Curtis distances (PERMANOVA, R² = 0.026, F = 19.418, *P* ≤ 0.001) and Jaccard distances (PERMANOVA, R² = 0.018, F = 13.121, *P* ≤ 0.001) revealed significant alterations in microbial community structure between atopic dermatitis (AD) and healthy controls, with inter-group variation exceeding intra-group variation (Fig. 4a, Supplementary Fig. 1a).

**Fig. 4.**
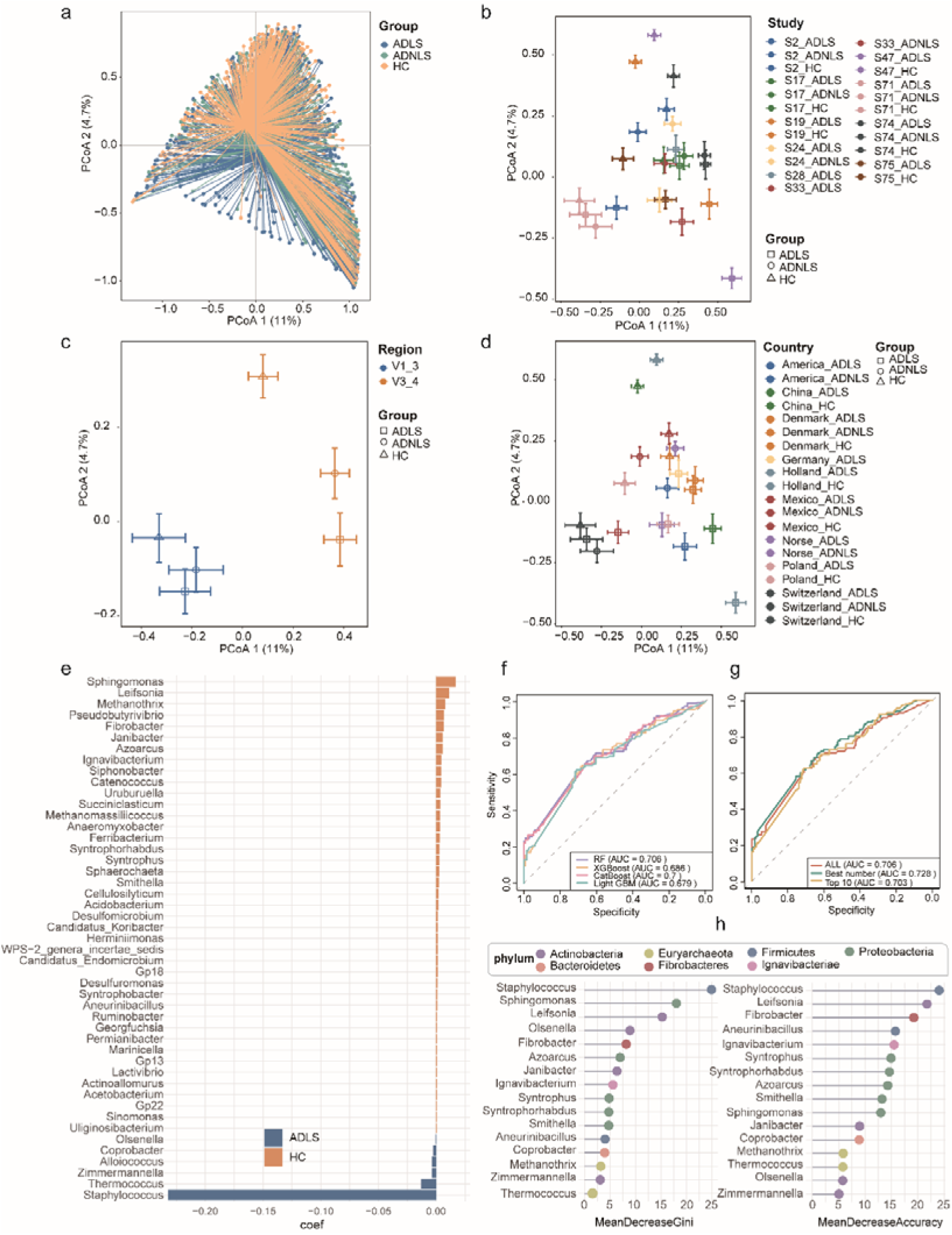
β diversity analysis and predictive model construction. **a** β-diversity analysis based on Bray-Curtis dissimilarity comparing differences between the ADLS, ADNLS, and healthy control (HC) groups. **b, c, d** Principal coordinate analysis (PCoA) based on Bray-Curtis dissimilarity showing the distribution of microbial community structures in the ADLS, ADNLS, and HC groups, grouped by different studies (b), variable regions (c), and countries (d). **e** Identification of the dominant taxa in the ADLS and HC groups using MMUPHin_MetaDA differential discriminant analysis (FDR ≤ 0.05). Blue represents dominant taxa in the ADLS group, while yellow represents dominant taxa in the HC group. **f** Random forest, XGBoost, CatBoost and LightGBM model constructed using all differential genera identified by MMUPHin_MetaDA, with the receiver operating characteristic (ROC) curve showing performance on the test set. **g** Random forest model constructed using differential genera identified by MMUPHin_MetaDA, with the receiver operating characteristic (ROC) curve showing performance on the test set. “ALL” represents the model constructed using all identified differential genera; “Best number” refers to the model based on the optimal number of differential genera selected by recursive feature elimination; “Top 10” represents the model based on the top 10 most important differential genera. **h** Evaluation of the relative importance of each genus in the predictive model based on mean decrease in Gini coefficient and accuracy. The color of the dots represents the phylum of each genus.

Given the potential influence of confounding factors such as cohort, variable regions, and geographical location on microbiome composition, we observed a reduction in batch effects after correction. However, significant differences persisted, prompting further subgroup analyses. The inter-group variation remained more pronounced than intra-group variation when stratified by cohort (PERMANOVA, R² = 0.0689, *P* ≤ 0.001), variable regions (PERMANOVA, R² = 0.007, F = 10.205, *P* ≤ 0.001), and country (PERMANOVA, R² = 0.066, F = 12.690, *P* ≤ 0.001) (Supplementary Table 1-1). The PCoA plots illustrating Bray–Curtis and Jaccard dissimilarities between the AD and control groups across different projects, variable regions, and countries are presented in Fig. 4b–d and Supp. Fig. 1b-d.

### Identification of skin microbial composite biomarkers for AD cross-cohorts

We employed the MMUPHin_MetaDA differential analysis method to identify the dominant bacterial taxa in the ADLS group compared to the HC group (FDR < 0.05). The results revealed that the AD group was enriched in Staphylococcus, Thermococcus, Zimmermannella, Alloiococcus, Coprobacter, and Olsenella (Fig. 4e, Supplementary Table 2). Next, we conducted a subgroup analysis based on different cohorts. Differentially abundant genera between the AD and HC groups were identified across studies S1 to S10 using three statistical methods: Permutation test, Wilcoxon rank-sum test, and LEfSe analysis (Supplementary Table 3-5). These differentially abundant genera (p < 0.05) were then used as features to examine their overlap across studies, as illustrated in Fig. 5a–c and Supplementary Fig. 2a-c. Additionally, Fig. 5d and Supplementary Fig. 2d presents the overlap of the combined differentially abundant genera identified by the three methods across different studies.

**Fig. 5.**
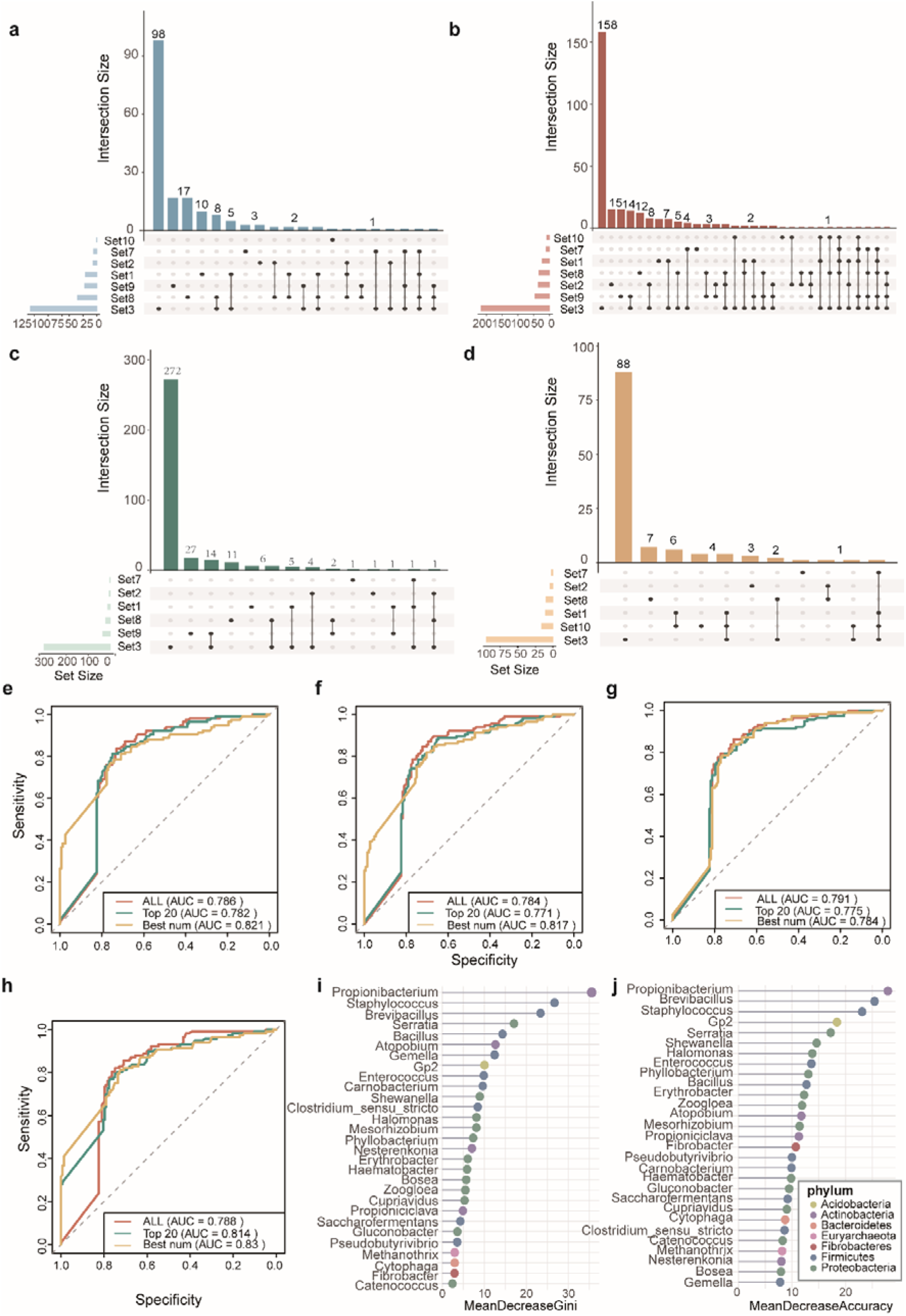
RF model was used to build a predictive model of genus-level abundant genera. **a–c** Overlapping differential genera between AD and healthy control (HC) groups across studies S1 to S10, as identified by Permutation testing (a), Wilcoxon rank-sum test (b), and the LEfSe algorithm (c). **d** Intersection of differential genera identified by all three methods within each study, highlighting shared taxa. **e–h** Receiver operating characteristic (ROC) curves of genus-level predictive models constructed using the Random Forest (RF) algorithm and tested on independent datasets. Models were built based on the union of differential genera identified by Permutation testing (e), Wilcoxon rank-sum test (f), and LEfSe (g) across all studies, as well as the intersection of genera identified by all three methods (h). **i–j** Relative importance of genera in the RF model (h), evaluated by mean decrease in accuracy (i) and Gini index (j).

To evaluate the diagnostic potential of microbiome signatures, we leveraged differentially abundant taxa identified across all dataset and within different subgroups to build predictive models using four machine learning algorithms: Random Forest (RF), XGBoost, LightGBM, and CatBoost. All samples were randomly partitioned into training (80%) and independent validation (20%) cohorts. Within the training set, microbial features derived from MMUPHin_MetaDA were used as inputs to train each of the four models to distinguish between atopic dermatitis (AD) and healthy control (HC) samples. Model performance was assessed on the validation set, yielding area under the receiver operating characteristic curve (AUC) values of 0.706 (RF), 0.686 (XGBoost), 0.699 (LightGBM), and 0.679 (CatBoost), respectively (Fig. 4f; Table 2). Among these, the Random Forest classifier exhibited the highest discriminatory power and was selected for subsequent analyses. To optimize predictive performance, we further constructed three RF classifiers based on distinct feature selection strategies: (1) the full set of taxa identified by MMUPHin_MetaDA, (2) an optimized subset selected through recursive feature elimination (RFE), and (3) the top 10 taxa ranked by feature importance. Corresponding AUCs in the validation cohort were 0.706, 0.728, and 0.703, respectively (Fig. 4g). The feature importance ranking of the best combination of differentially abundant taxa as classification features is illustrated in Fig. 4h.

**TABLE 2.**
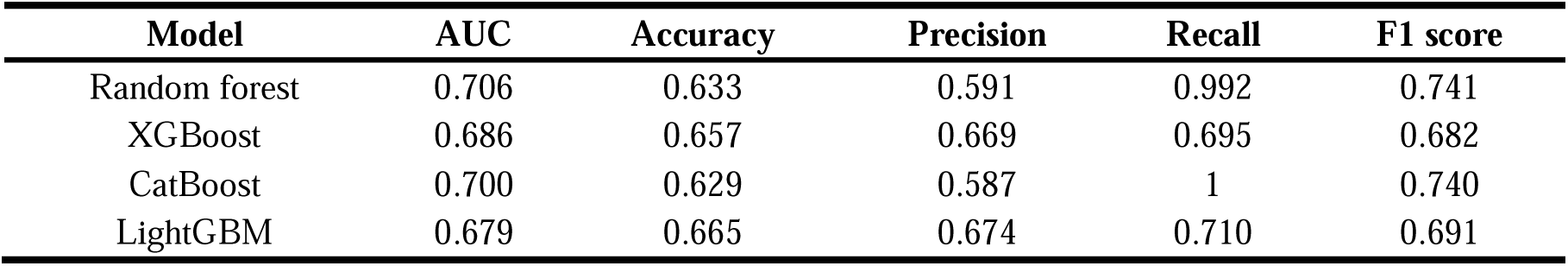
Predictive performance of four machine learning models for Alzheimer’s disease classification.

Furthermore, in the subgroup analyses, we constructed random forest models on the training set using the union of differentially abundant taxa identified by the three differential analysis methods for each project, as well as the best combinations of differentially abundant taxa determined by recursive feature elimination, and the top 20 taxa based on feature importance ranking. Based on Permutation, the predicted AUCs were 0.786, 0.821, and 0.782 (Fig. 5e); based on Wilcoxon rank-sum test, the predicted AUCs were 0.784, 0.817, and 0.771 (Fig. 5f); and based on LEfSe, the predicted AUCs were 0.791, 0.784, and 0.775 (Fig. 5g). Additionally, when the intersection of differentially abundant taxa obtained from the three methods was used as classification features, random forest models were constructed, and the predicted AUCs were 0.788, 0.83, and 0.814 (Fig. 5h). The feature importance rankings of the best combinations of differentially abundant taxa as classification features are shown in Fig. 5i and 5j.

## MATERIALS AND METHODS

### Study search, selection, and inclusion

We conducted a search in the PubMed database. The search strategy and search terms are constructed as follows: ((((((((((((((((Dermatitis, Atopic[MeSH Terms]) OR (Atopic Dermatitides[Title/Abstract])) OR (Atopic Dermatitis[Title/Abstract])) OR (Dermatitides, Atopic[Title/Abstract])) OR (Neurodermatitis, Atopic[Title/Abstract])) OR (Atopic Neurodermatitides[Title/Abstract])) OR (Atopic Neurodermatitis[Title/Abstract])) OR (Neurodermatitides, Atopic[Title/Abstract])) OR (Neurodermatitis, Disseminated[Title/Abstract])) OR (Disseminated Neurodermatitis[Title/Abstract])) OR (Neurodermatitides, Disseminated[Title/Abstract])) OR (Eczema, Atopic[Title/Abstract])) OR (Atopic Eczema[Title/Abstract])) OR (Eczema, Infantile[Title/Abstract])) OR (Infantile Eczema[Title/Abstract])) AND (((((((((((((((((Microbiota[MeSH Terms]) OR (Microbiotas[Title/Abstract])) OR (Microbial Community[Title/Abstract])) OR (Community, Microbial[Title/Abstract])) OR (Microbial Communities[Title/Abstract])) OR (Microbial Community Composition[Title/Abstract])) OR (Community Composition, Microbial[Title/Abstract])) OR (Composition, Microbial Community[Title/Abstract])) OR (Microbial Community Compositions[Title/Abstract])) OR (Microbial Community Structure[Title/Abstract])) OR (Community Structure, Microbial[Title/Abstract])) OR (Microbial Community Structures[Title/Abstract])) OR (Microbiome[Title/Abstract])) OR (Microbiomes[Title/Abstract])) OR (Human Microbiome[Title/Abstract])) OR (Human Microbiomes[Title/Abstract])) OR (Microbiome, Human[Title/Abstract]))) AND ((((((RNA, Ribosomal, 16S[MeSH Terms]) OR (16S rRNA[Title/Abstract])) OR (rRNA, 16S[Title/Abstract])) OR (16S Ribosomal RNA[Title/Abstract])) OR (RNA, 16S Ribosomal[Title/Abstract])) OR (Ribosomal RNA, 16S[Title/Abstract])). The study was limited to research published from the inception of the database up until May 1, 2025. Furthermore, the search was restricted to studies without language limitations.

### Data set collection

Raw sequence data and metadata were retrieved from the NCBI Short Read Archive (SRA), European Nucleotide Archive (ENA), or the article content. Subsequently, samples were excluded if metadata were unavailable. The study information for the selected studies included demographic details and methods such as recruitment area, sample size, average age, diagnostic methods for AD, PCR primers, sequencing platform, hypervariable region sequences, and whether raw data were recorded. Prior to downstream analysis, samples were grouped accordingly.

### Processing of 16SrRNA gene sequences

We processed the raw sequencing reads of the 16S rRNA gene sequences using QIIME2^49^ and USEARCH^50^ pipelines^51^. When paired-end sequence files were provided by the study, the fastq_mergepairs command was used to join the reads. The fastq_minmergelen 250 command filtered out reads shorter than 250 bp, and the merged reads were filtered using a maximum error rate of 1%. Ultimately, 84.2% of the raw reads were retained. Representative sequences were selected from at least 6 completely identical sequences. Taxonomy was assigned using the RDP^52^ database (version 2.14, 2024) with the Bayesian classifier, and operational taxonomic units (OTUs) were generated using UPARSE, which sorted 6,256,417 quality-filtered sequences into 18,162 OTUs with 97% sequence homology. Additionally, when calculating α- and β-diversity, the sequencing depth was set to 11,226.

### Confounder analysis and removing batch effect

To evaluate the influence of confounding factors, including cohort, sequencing region, country, and sampling site, on the skin microbiome and to compare their relative effects to atopic dermatitis (AD) disease status, we performed an ANOVA-type analysis^53^. This approach enables the assessment of whether disease status, as well as each confounding factor, significantly alters the abundance of specific microorganisms. Furthermore, it quantifies the variance in microbial abundance explained by both disease status and these confounding variables. Ultimately, this analysis provides insights into the factors that may shape the microbiome composition and determines the extent to which their influence surpasses the effect of AD itself on the skin microbiome.

If significant effects of confounding factors were observed, we used the “adjust_batch” function from the R package wrapped in MMUPHin^36^ (Meta-Analysis Methods with a Uniform Pipeline for Heterogeneity in microbiome studies) to remove batch effects. Subsequently, a meta-analysis of differential abundance was conducted on the batch-corrected microbial abundance matrix using the “lm_meta” command.

### Microbial community ecological profiling

The Shannon diversity index^54^, Pielou’s evenness index^55^, and Simpson diversity index^56^ were used to compare the α-diversity differences between the lesion and non-lesion skin regions of AD patients and healthy controls across different studies. Meta-analysis was performed using the R package “metafor“^57^. For continuous variables, the weighted standardized mean difference (SMD) and 95% confidence intervals (CIs) were calculated to compare the two groups, with Hedges’ g chosen as the final effect size. A p-value of less than 0.05 was considered statistically significant. To assess the extent of heterogeneity across studies, we used Cochran’s Q test and I² statistics. Given the anticipated high degree of heterogeneity, pooled estimates were derived using random-effects models.

Beta diversity between samples was assessed using Bray–Curtis dissimilarity^58^ and Jaccard dissimilarity^59^, and visualized through principal coordinate analysis (PCoA)^60^. Similarity analysis (ANOSIM)^61^ was performed based on Bray–Curtis dissimilarity to test whether the differences between groups were significantly greater than the within-group differences. We also employed permutational multivariate analysis of variance (PERMANOVA)^62^ to evaluate the microbial differences between the lesion region, non-lesion region of AD patients, and healthy controls.

### Differential abundance analysis to identify skin microbes

To better identify microbial biomarkers associated with AD, we employed three classic microbiome differential analysis methods: linear discriminant analysis (LDA) effect size (LEfSe)^39^, permutation test^63^, and Wilcoxon rank-sum test^38^. In the sub-cohort analysis, these three methods were used to perform differential testing between the ADLS, ADNLS, and HC groups for each of the 10 datasets, identifying significantly different microbial taxa, and their union was selected. The intersection of the differentially abundant taxa identified by the three methods was then used as biomarkers for subsequent analysis. Additionally, for the cross-cohort overall analysis, differential abundance testing was performed using the “lm_meta” function from the aforementioned R package “MMUPHin”.

### Feature selection and classifier model building and performance based on **machine-learning**

To determine whether composite microbial biomarkers can distinguish AD patients from the control group, a random forest^64^, XGboost^65^, CatBoost^66^, LightGBM^67^ classification model was constructed using the relative abundance of microorganisms identified through differential analysis. Five-fold cross-validation was performed using the “caret^68^” and “randomForest^69^”, “xgboost^70^”, “catboost”, “lightgbm” packages in R (ver. 4.0.3, R Foundation for Statistical Computing)^64^. During model construction, the 10 cross-cohort amplicon sequencing datasets were first divided into five parts. Four parts were randomly selected for model training, while the remaining independent part (test data) was used for model validation. Subsequently, the model performance and the clinical diagnostic capability of the microbial biomarkers were assessed using the receiver operating characteristic (ROC) curve, along with the area under the curve (AUC), accuracy, precision, recall, and F1 score metrics. The importance of each feature in the model was evaluated and ranked based on mean decreasing accuracy and the Gini coefficient. Recursive feature elimination^71^ was then applied to select the optimal number of features, determining the best-performing model. Additionally, features were ranked by importance, and models were built with varying numbers of features to identify the most streamlined model.

## DISCUSSION

This study represents the first comprehensive meta-analysis of skin microbiome 16S rRNA sequencing data in AD patients. While previous studies have demonstrated a strong association between dysbiosis of the skin microbiome and AD pathogenesis, two key questions remain unresolved: the reproducibility of microbial biomarkers identified from skin samples across different cohorts and populations, and the reliability and accuracy of these biomarkers as diagnostic indicators. By conducting a cross-cohort integrative analysis (CCIA) of raw sequencing data and metadata from 10 studies, encompassing 1,522 samples, we identified consistent characteristics of the commensal skin microbiome, examined alterations in the skin microbiota of AD patients, and evaluated their diagnostic potential as biomarkers. Our findings provide novel insights into the role of the skin microbiome in AD pathophysiology.

We found significant differences in both α-diversity and community composition of the skin microbiome between the AD and control groups. Specifically, the α-diversity of the skin microbiome was significantly reduced in AD patients, consistent with previous studies, suggesting that decreased microbial diversity may be involved in AD pathogenesis. Additionally, we observed no significant differences in microbial composition between lesion and non-lesion skin in AD patients, which may indicate systemic alterations in the skin microbiome. This finding suggests that AD pathogenesis could involve a broader dysbiosis beyond localized skin lesions^8,72^. These results highlight the potential role of the skin microbiome in the early diagnosis and preemptive intervention of AD.

The structure and function of the skin microbiome are influenced by various confounding factors, such as cohort differences, sequencing variable regions, geographical location, and sampling sites. To minimize the impact of these confounders, we applied the MMUPHin tool to correct for batch effects. Although batch correction reduced the impact on the skin microbiome composition, significant effects remained. Therefore, we further conducted subgroup analyses stratified by different batches to account for these variations.

Using Permutation tests, the Wilcoxon rank-sum test, and Linear Discriminant Analysis Effect Size (LEfSe), we identified a set of microbial biomarkers associated with AD. These biomarkers exhibited significant differences between AD patients and healthy controls, demonstrating high diagnostic value. The predictive model constructed using machine learning algorithms achieved robust performance in the validation cohort (AUROC: 0.83), further supporting the potential of these microbial biomarkers for AD diagnosis^3,73^.

Additionally, we observed that across both cross-cohort analyses and batch-specific subgroup analyses, Staphylococcus was consistently identified as a differentially abundant genus by multiple analytical approaches. The abundance of Staphylococcus was significantly higher in the skin of AD patients compared to healthy controls, consistent with previous findings. Notably, Staphylococcus aureus was significantly enriched in AD skin and was associated with the expression of disease-related genes involved in skin barrier function, immune activation, and tryptophan metabolism^27^. Furthermore, the increased abundance of Staphylococcus capitis and Staphylococcus lugdunensis in AD skin correlated with disease severity^40^.

The skin microbiome not only influences barrier function and immune responses but may also interact with the host’s systemic physiology through neural and endocrine pathways. For instance, certain microbes may exacerbate AD symptoms by producing metabolic byproducts that modulate skin inflammation. Moreover, alterations in the skin microbiome could be associated with impaired host immune regulation, further contributing to the onset and progression of AD^15,74^.

### Strengths and limitations

Despite the meaningful findings of this study, several limitations should be acknowledged. First, variations in sample selection, sequencing platforms, and data analysis methods across studies may affect the accuracy and comparability of the results. Future research should adopt more standardized and uniform approaches for data collection and analysis to minimize these discrepancies^75^. Second, while this study primarily focused on the composition and function of the skin microbiome, the causal relationship between microbial alterations and host responses remains unclear. Further experimental studies, including animal models and interventional research, are needed to validate these associations and explore the underlying molecular mechanisms^76,77^. Additionally, considering the dynamic nature of the skin microbiome, future studies should investigate microbial changes across different stages of AD, as well as their associations with disease progression and treatment response^78,79^.

## Conclusion

In summary, this study integrates data from multiple cohorts to reveal both the common and distinct alterations of the skin microbiome in AD. These findings provide new insights into the role of the skin microbiome in AD, shedding light on host-microbiome interactions, early diagnosis, and targeted interventions for atopic dermatitis. Future research should further explore the mechanistic interactions between the skin microbiome and the host to develop microbiome-based diagnostic and therapeutic strategies for AD.

## DATA AVAILABILITY

This study incorporates data from previously published research. The data were obtained from NCBI Short Read Archive (SRA) and European Nucleotide Archive (ENA), with accession numbers listed in the Table 1.

## COMPETING INTERESTS

The authors declare that the research was conducted in the absence of any commercial or financial relationships that could be construed as a potential conflict of interest.

## AUTHORS’ CONTRIBUTIONS

All authors contributed to the pipeline development and workflow of analyses. The initial idea and framework were conceived by Yihua Wang. Yihua Wang contributed to the pipeline test and analyses. This protocol was written by Yihua Wang and Bingqiang Liu, and revised with from Bingqiang Liu’ suggestions.

## Notes

### Competing Interest Statement

The authors have declared no competing interest.

